# Computational analysis of 4-1BB-induced NFκB signaling suggests improvements to CAR cell design

**DOI:** 10.1101/2022.04.27.489659

**Authors:** Vardges Tserunyan, Stacey D. Finley

## Abstract

**Background:** Chimeric antigen receptor (CAR)-expressing cells are a powerful modality of adoptive cell therapy against cancer. The strength and dynamics of signaling events initiated upon antigen binding depend on the costimulatory domain within the structure of the CAR. One such costimulatory domain is 4-1BB, which affects cellular response via the NFκB pathway. However, the quantitative aspects of 4-1BB-induced NFκB signaling are not fully understood.

**Methods:** We developed an ordinary differential equation-based mathematical model representing canonical NFκB signaling activated by CAR-4-1BB. We first performed a global sensitivity analysis on our model to quantify the impact of kinetic parameters and initial protein concentrations. We ran Monte Carlo simulations of cell population-wide variability in NFκB signaling and used Kraskov’s algorithm to quantify the mutual information between the extracellular signal and different levels of the NFκB signal transduction pathway.

**Results:** We determined that in response to a wide range of antigen concentrations, the magnitude of the transient peak in the nuclear concentration of NFκB varies significantly, while the timing of this peak is relatively consistent. Our global sensitivity analysis showed that the model is robust to variations in parameter values, and thus, its quantitative predictions would remain applicable to a broad range of parameter values. Next, we found that overexpressing NEMO and disabling IKKβ deactivation are predicted to increase the mutual information between antigen levels and NFκB activation.

**Conclusions:** Our modeling predictions provide actionable insights to guide CAR development. Particularly, we propose specific manipulations to the NFκB signal transduction pathway that can fine-tune the response of CAR-4-1BB cells to the antigen concentrations they are likely to encounter.

## Background

In the last two decades, adoptive cell therapy has emerged as a powerful tool for combatting cancer by enhancing the immune response against cancer cells (1). One successful approach is to confer adoptive immune cells with specificity against antigens expressed on cancer cells by engineering immune cells to express chimeric antigen receptors (CAR) (2). Currently, T cells are the predominant immune cell type used for CAR-based immunotherapy, and multiple CAR-T cell therapies have received FDA approval (3). The key step in this approach is to produce T cells expressing a CAR that activates the T cell upon encountering the target antigen. CARs are expressed on the cell surface and feature an antigen recognition domain derived from single-chain variable fragments of monoclonal antibodies, while their cytoplasmic portion includes different combinations of intracellular signaling domains. The first generation of CARs had a single cytoplasmic CD3ζ signaling domain connected to the recognition domain by a transmembrane linker. Later advances resulted in second-generation CARs, which have one additional costimulatory signaling domain originating from signaling endodomains of natively occurring costimulatory receptors (4). 4-1BB (also referred to as CD137) is one such costimulatory domain frequently incorporated in the structure of the CAR (5).

The effects mediated by the 4-1BB domain in immune cells have been tracked to the Nuclear Factor κB (NFκB) signaling pathway (6–8), which is comprised of two branches, termed canonical and non-canonical NFκB signaling (9,10). The canonical NFκB signaling is triggered in a matter of minutes after stimulation, while the non-canonical system is slower and depends on protein translation to exert its effect (11). In addition, there is a functional difference between the two branches: the sets of genes activated by each are not identical (12). The genes of many cytokines, such as those of IL-1α/β (13,14), IL-6 (15), IL-10 (16) and IFNγ (17), as well as perforin (18), show experimental evidence of canonical NFκB regulation. Meanwhile, the non-canonical NFκB pathway was found to be responsible for improved survival rates among 4-1BB-bearing second-generation CAR-T cells (8). Thus, investigating NFκB signaling initiated by 4-1BB can help gain a deeper understanding of the observed biological features of CAR cells.

Computational systems biology provides powerful tools with which we can explore signaling systems without the need for expensive and prolonged experimentation. For example, this approach was successful in describing T cell receptor-induced activation of MAPK/ERK signaling (19). Recently, our research group successfully utilized an ordinary differential equation (ODE)-based model to describe MAPK signaling in a heterogeneous CAR-T cell population expressing the second generation CD3ζ-CD28 construct and contrasted them with the CAR-T cells lacking the CD28 domain (20,21). Several mathematical models have been published for NFκB signaling as well. Some of these mathematical models have mostly focused on downstream events of the pathway, replacing the dynamics of IKKβ with an assumed activation profile (22–24), while other models have incorporated a more explicit account of upstream processes (25–27). Mathematical modeling was also used to analyze signal transduction capabilities of the NFκB pathway used by viewing it as an information transmission channel. This work has shown that the pathway can encode approximately 1 bit of information at most, i.e., whether the extracellular stimulus is present or absent (28,29). Insights from the information-theoretic analysis of NFκB signal transmission were extended to the expression patterns of genes affected by nuclear levels of NFκB (30). However, none of these analyses addresses NFκB signaling in the context of cells expressing CAR receptors. In addition, while some published models include aspects of NFκB activation via 4-1BB (31), they are based on a logic-gate approach and do not capture the dynamic chemical interactions occurring in response to stimulation.

Hence, in our current work, we set out to develop a mechanistic ODE-based model of canonical NFκB signaling initiated by antigen binding to CAR with a 4-1BB co-stimulatory domain. We applied the model to quantitatively investigate the features of 4-1BB-mediated NFκB signaling. We found that in response to antigen binding, 4-1BB initiates a transient increase in the nuclear concentration of NFκB. The timing of the peak NFκB concentration is relatively consistent over a broad range of antigen concentrations, while its magnitude shows a much greater dependence on the antigen level. Next, we found that very few of the model parameters have a significant impact on the activation time and magnitude of the NFκB pathway, indicating that the behavior of the model would be robust to large variations in kinetic parameters and protein concentrations. Finally, by analyzing the mutual information between antigen concentration and successive levels of the NFκB pathway in conditions of simulated biochemical variability, we proposed potential improvements to CAR-4-1BB cells. Specifically, our results suggest that the overexpression NEMO or the disabling of IKKβ deactivation would have the opposite effect. Altogether, we provide a quantitative framework that can be used to guide the development and optimization of 4-1BB-induced CAR signaling.

## Methods

### Model Structure

By performing an extensive literature search, we compiled a list of key protein-protein interactions and chemical reactions, which form the core of the activation of the canonical NFκB pathway. In our model, in response to antigen binding, the 4-1BB domain of the CAR induces the formation of a signalosome, in which TRAF2 trimers play a key role (7,32,33). Acting as a ubiquitin-ligase, TRAF2 promotes the attachment of long K63 ubiquitin chains to RIP1 (34,35). Once RIP1 is ubiquitinated, its K63 ubiquitin chains enable the docking of TAB1/2/3, the adaptor subunits of the TAK enzymatic complex (36). Similarly, the IKK multimeric complex is docked with NEMO (also known as IKKγ) as an adaptor subunit to the K63 ubiquitin chains of RIP1 (37). The protein TAK1, the enzymatic subunit of the TAK complex, phosphorylates the β subunit of the IKK complex in a ubiquitin-dependent manner (38). IKKβ activation is transitory since phosphorylation of a cluster of 10 serine residues at a regulatory domain deactivates it (39,40). Catalytically active IKKβ, unlike the subunit IKKα, phosphorylates the protein IκBα, targeting it for proteasomal degradation (10,41,42). In resting cells, IκBα tightly associates with cytoplasmic NFκB, preventing it from shuttling to the nucleus. However, the degradation of IκBα under the influence of catalytically active IKKβ releases cytoplasmic NFκB from this sequestration and makes it free to translocate to the nucleus, where it can function as a transcription factor and alter the expression profile of the cell (9).

These reactions were represented as interaction rules in RuleBender (v2.3.1) (43). Then, RuleBender was used to generate a corresponding set of ordinary differential equations (ODEs) describing the temporal dynamics of the system. MATLAB (v2019a/b) was used to integrate the system of ODEs to obtain time courses of the species involved. Kinetic parameters and protein concentrations were compiled from prior models of different components of the system or direct experimental measurements, with a detailed list provided in the Supplemental File.

### Activation profiles and dose response curves

We simulate binding of CD19, a common target for CAR therapies, to the antigen recognition domain of the CAR. Prior experimental measurements of the concentration of CD19 estimated its surface concentration to have a mean of 3.4 molecules per µm^2^ on NALM-6, a human acute lymphoblastic leukemia-derived cell line, while the CD19 concentration on primary myeloma cells was found to be in the range of 0.16-5.2 µm^2^ (44). We took these values as a basis for our simulations of activation profiles and dose response curves but used the wider range of 10^−0.5^-10^2.5^ molecules per µm^2^ in order to gain a complete picture of dynamic behaviors captured by the model in a broader range of antigen concentrations.

### Sensitivity analysis

We performed sensitivity analysis to determine how strongly model inputs (kinetic parameters and initial protein concentrations) affect the quantitative predictions of the model. We use the Extended Fourier Amplitude (eFAST) method of quantifying global sensitivity of a mathematical model to underlying parameter values (45). The sensitivity coefficients are obtained by simultaneously varying parameters of interest at different pre-assigned signature frequencies and evaluating system output for each subsequent value of the parameter vector. Then, the Fourier transform of sequential values of system output is taken to reveal its spectral composition. The underlying assumption is that if a parameter influences system output to a significant extent, then its periodic variation according to a certain signature frequency would cause large variation in model output at that frequency, as revealed by a higher amplitude in the Fourier transform. The significance of different parameters is evaluated in comparison with the sensitivity coefficient attained by a “dummy” parameter, a parameter with no involvement in the system at all. Following the spectral decomposition of the output, if the amplitude at any system parameter’s characteristic frequency is lower than that for the “dummy” parameter, the system parameter is taken to have no significant impact on system output.

### Monte Carlo simulations

We considered a population of CAR-expressing cells by accounting for cell-to-cell variability in protein concentrations. Motivated by prior experimental findings (46), we modeled the intrinsic noise of the population response by assuming that proteins involved in signal transduction in each cell have varying initial concentrations subject to a lognormal distribution. We chose the location parameter of each distribution so that the median concentration across the CAR cell population would equal the accepted value of the protein concentration. The scale parameter was taken to be the same for all proteins, and different hypothesized values of this scale parameter were investigated to gain a more complete picture (termed “noise parameter” throughout). We included variability in target antigen concentration by assuming a lognormal distribution and choosing the location and scale parameters such that the distribution would mostly lie within the measured range of 0.16-5.2 µm^2^ for myeloma cells and have a mean of approximately 3.4 µm^2^ as measured for lymphoma cells (44). This random sampling with subsequent simulation was repeated 2000 times to yield representative distributions.

### Mutual information

Mutual information is a quantity measuring the mutual dependence of two random variables (47). As such, its value is zero if and only if two random variables are independently distributed, and a positive number if they depend on each other. We used the Kraskov algorithm for computing mutual information based on distribution entropies, with the entropies estimated through a *K*-nearest neighbors procedure (48).

We computed the mutual information between antigen levels and five different stages of signal transduction: concentration of antigen-bound CAR, amount of TRAF2 bound to the 4-1BB domain, amount of RIP1 bound to the TRAF2 signalosome, concentration of enzymatically active TAB2/TAK1, peak concentration of enzymatically active NEMO/IKKβ, and peak nuclear concentration of NFκB. In order to quantify the amount of information reaching a certain level along signal transduction, we randomized the concentrations of all proteins upstream of that level, leaving the rest of the protein concentrations unchanged. Then, we stimulated the system by a randomly sampled antigen concentration. Since only the steps leading up to the given level were randomized, any variation at this level would reflect upstream stochasticity and not an artifact of the simulation. On the other hand, by randomizing all upstream proteins, we ensured that the contribution of every upstream component to overall stochasticity is accounted for. By performing this procedure for five different levels of the signal transduction pathway, we were able to track how much mutual information is shared between the antigen concentration and a given level of the signaling pathway.

## Results

### Model structure and dose response

In order to obtain a quantitative description of signaling events triggered by antigen binding CAR-4-1BB cells, we developed an ODE-based mechanistic model (Fig. 1, Supplemental File). The biochemical reactions included in the model are based on prior experimental and modeling studies available in literature. Given the paucity of quantitative experimental measurements for 4-1BB in immune cells, the model is largely based on data from studies performed using fibroblasts. We utilize this model to investigate various aspects of 4-1BB signaling and generate novel hypotheses about how to enhance the signaling promoted by this co-stimulatory domain.

**Figure 1:**
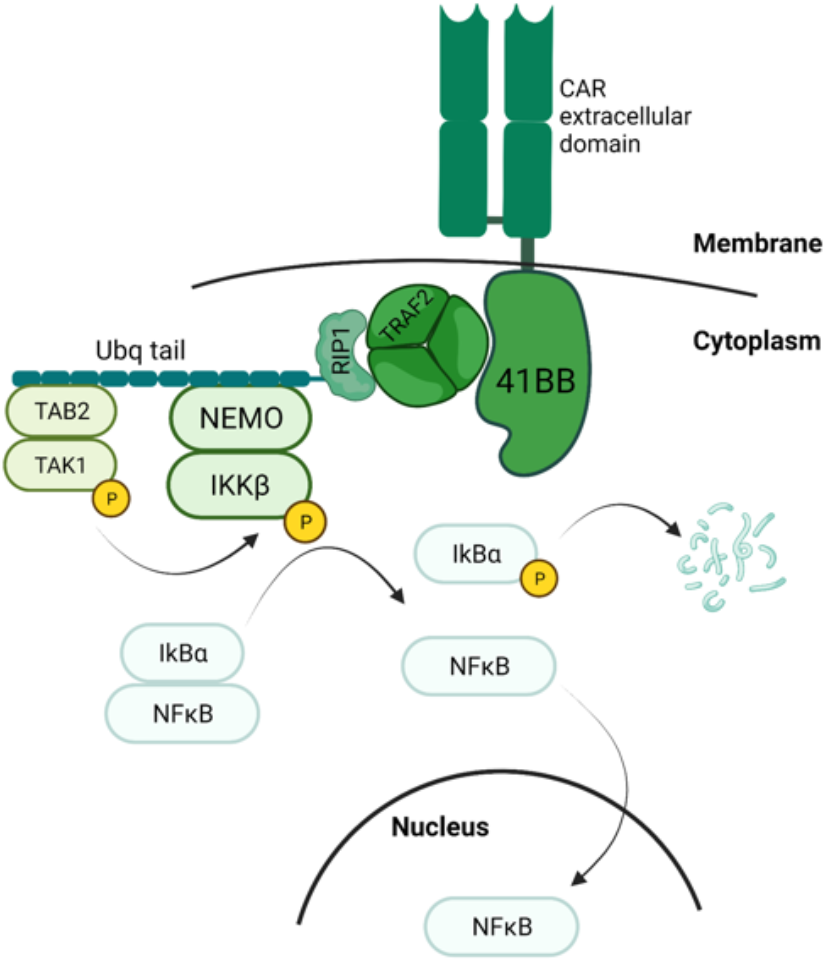
A schematic outline of events leading up to the nuclear translocation of canonical NFκB.

To understand the signaling dynamics predicted by our model, we generated time courses for enzymatically active IKKβ, which is a key convergence point for NFκB-activating signals (49–51), and nuclear NFκB for a range of physiologically relevant antigen concentrations values (Fig. 2). The model predicts that in response to persistent antigen binding, there is only transient activation of IKKβ. The IKKβ concentration is predicted to exhibit a rapid-onset, short-term peak typically occurring 10-15 minutes after stimulation (Fig. 2A). We observed a similar transient peak for the nuclear concentration of NFκB. Here, the peak was broader, with the maximum concentration occurring after approximately 20-40 minutes of stimulation with physiologically relevant antigen concentrations (Fig. 2B).

**Figure 2:**
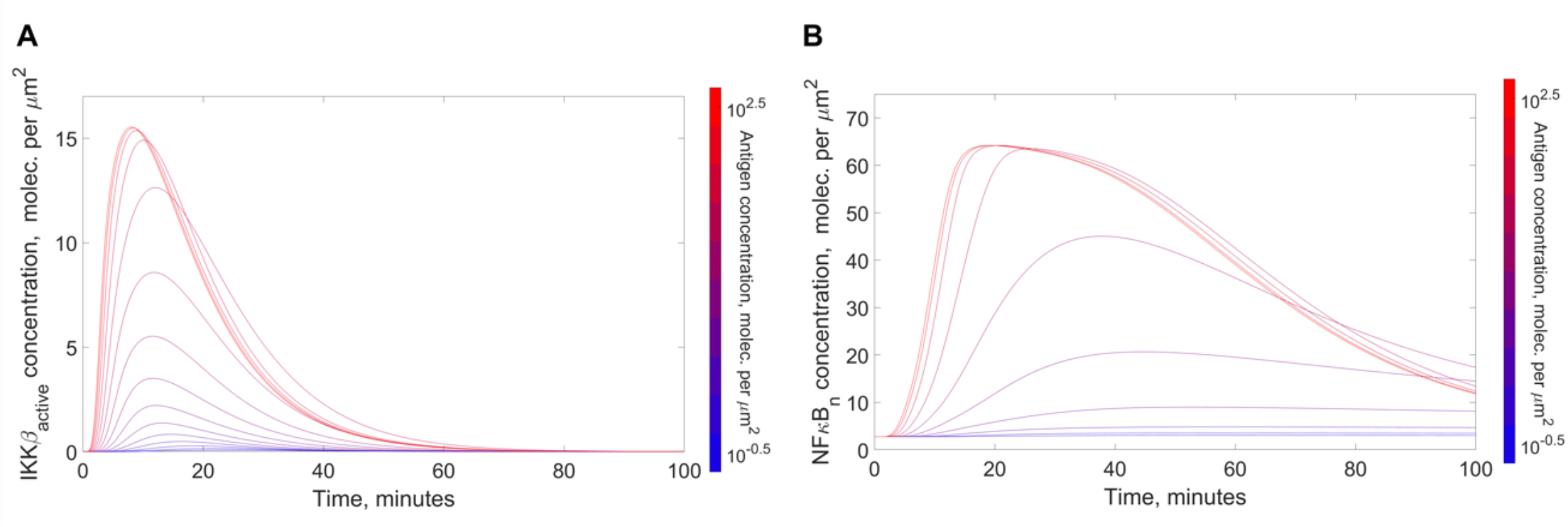
Activation profiles for IKKβ and NFκB in response to antigen binding the CAR. The pathway was stimulated with 10 different antigen concentrations *in silico* and activation profiles for IKKβ and NFκB were recorded for each antigen concentration. (A) Enzymatically active IKKβ; (B) Nuclear concentration of NFκB.

Next, we obtained dose response curves for the activation of IKKβ and NFκB (Fig. 3). Notably, only a small fraction of the total IKKβ pool (< 15%) was activated even at its maximal level and with saturating antigen levels (Fig. 3A). Peak concentration achieved by active IKKβ had a sigmoidal dependence on antigen concentration and was sensitive to antigen levels. A 1000-fold change in antigen concentration results in a 193-fold change in active IKKβ peak concentration. In contrast, the timing of the IKKβ peak was relatively robust to the antigen concentration. Specifically, a 1000-fold range of antigen concentrations produces only a 3-fold change in the time at which IKKβ peaked (Fig. 3B). The nuclear concentration achieved by NFκB increased with increasing antigen concentration (21-fold change in the response range) and showed a sigmoidal dependence on antigen concentration (Fig. 3C). In addition, the timing of peak nuclear NFκB concentration was relatively consistent for different antigen concentrations (2.9-fold change in the response range, Fig. 3D).

**Figure 3:**
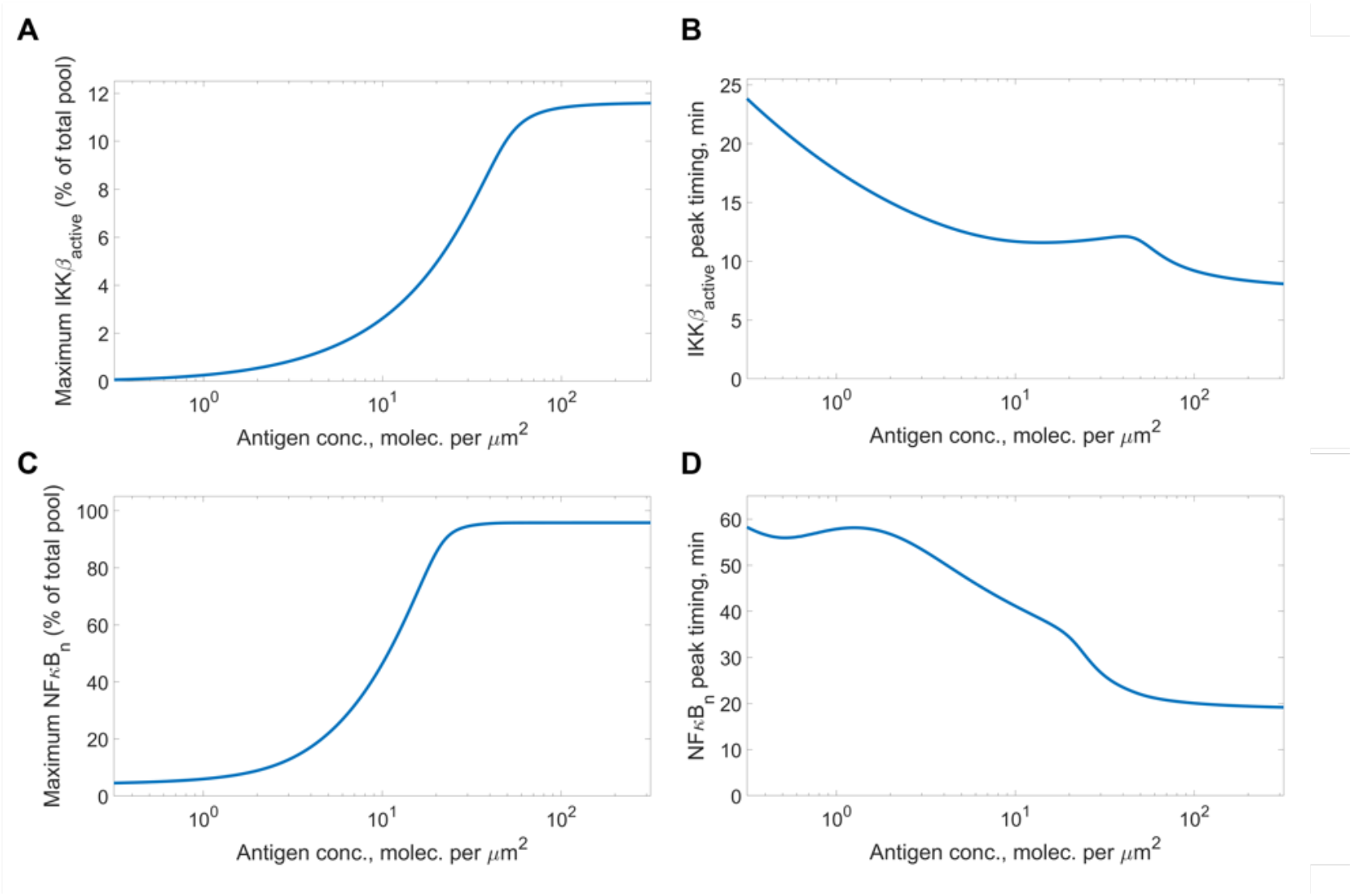
Dose response curves for IKKβ and NFκB in response to antigen binding the CAR. Relative magnitude and timing of the peak concentration of active IKKβ and nuclear NFκB were recorded for a range of antigen concentrations. (A) Relative magnitude of the peak concentration of enzymatically active IKKβ; (B) Relative magnitude of the peak concentration of nuclear NFκB; (C) Timing of the peak concentration of enzymatically active IKKβ; (D) Timing of the peak concentration of nuclear NFκB.

### Sensitivity of the output to kinetic parameters and protein concentrations

Once we established basic features of pathway activation predicted by our model, we proceeded to evaluate how the model output is impacted by the model parameters. This would allow us to see how sensitive the quantitative predictions from our model are with respect to choices of specific values for kinetic constants and protein initial concentrations. To this end, we used the Extended Fourier Amplitude Sensitivity Test (eFAST), a method of global sensitivity analysis, on the kinetic constants for regimes of low, medium and high antigen stimulation (Fig. 4A). We evaluated the impact of the kinetic constants on four measures of the system output: (1) the timing of peak active IKKβ concentration, (2) the magnitude of peak active IKKβ concentration, (3) the timing of peak nuclear NFκB concentration and (4) the magnitude of peak nuclear NFκB concentration.

**Figure 4:**
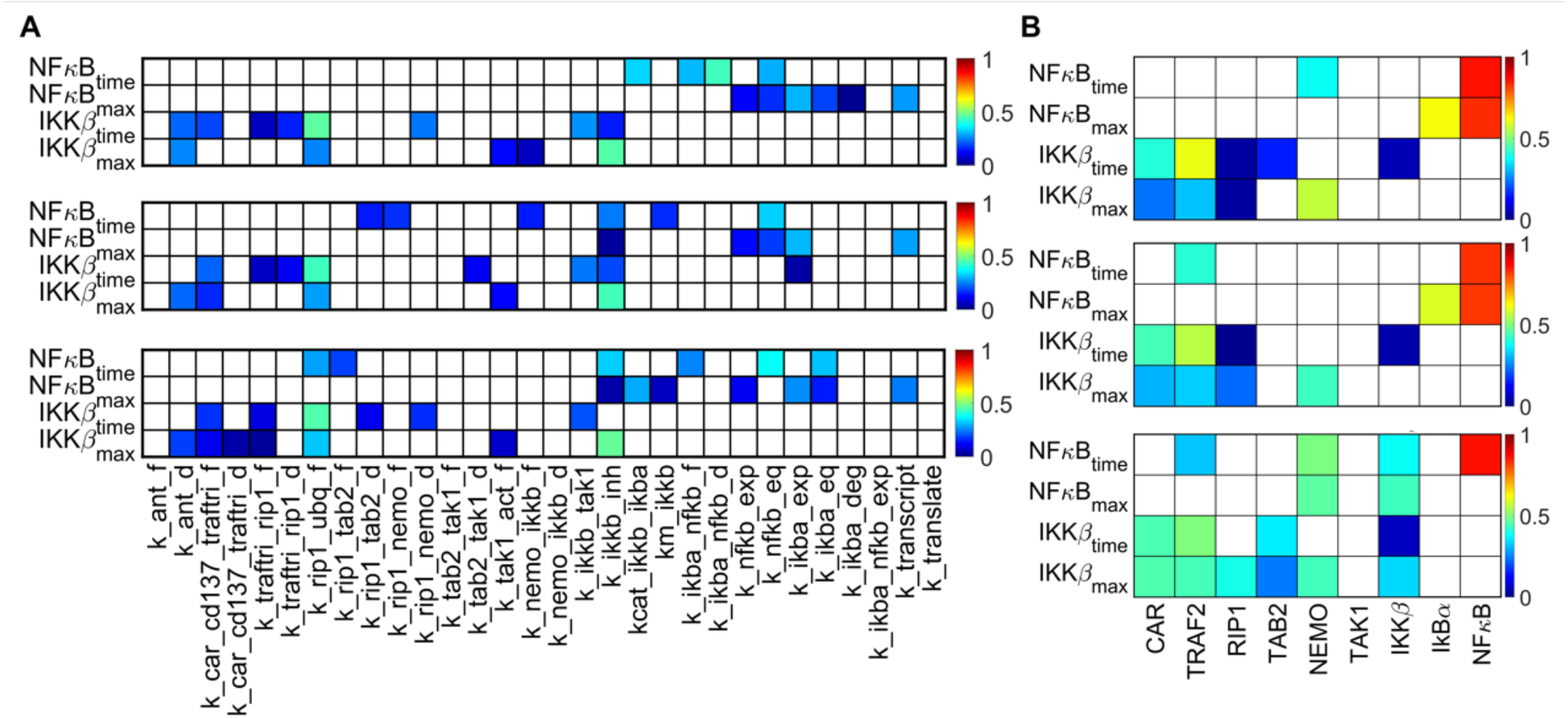
eFAST total sensitivity indices of model output with respect to model parameters. (A) Sensitivity indices for kinetic parameters; (B) Sensitivity indices for protein initial concentrations. From top to bottom: low (0.10 molec. per µm^2^), medium (1.0 molec. per µm^2^) and high (10 molec. per µm^2^) antigen concentration.

Based on our results, the responses of both IKKβ and NFκB appear to be relatively robust with respect to variations in the kinetic parameters. In fact, only very few parameters are shown to significantly impact the timing and magnitude of their peak concentrations. For example, at most eight of the 30 kinetic constants influence the magnitude or timing of nuclear NFκB across the range of antigen concentrations we considered (considering a sensitivity index > 0.3). Particularly, at low antigen concentrations, nuclear import/export rates of NFκB and IkBα strongly influence peak nuclear NFκB concentration. At higher antigen concentrations, parameters related to the enzymatic activity of IKKβ exert more influence.

Next, we performed eFAST on the initial concentrations of the proteins involved in signal transduction (Fig. 4B). We observed a very strong dependence of the timing and magnitude of peak nuclear NFκB concentration on the total cellular amount of NFκB. In addition, with non-saturating antigen concentrations, the NFκB response is predicted to depend on the amount of IkBα, the protein which sequesters NFκB in the cytoplasm. Interestingly, while the activation of the upstream protein IKKβ depends on the amounts of many of the upstream proteins, those upstream species do not significantly affect NFκB.

The robustness of model output to kinetic parameters and initial concentrations of pathway proteins demonstrates that the dynamics of the NFκB pathway predicted by the model would remain essentially similar across a wide range of parameter values. This indicates that the quantitative insights derived from the model would be applicable to a broad range of systems, including engineered CAR-4-1BB cells, even though the model is based on studies performed on fibroblasts. This provides confidence that we can gain biological insights form the model predictions.

### Mutual information carried by the NFκB pathway about antigen concentration

Signal transduction pathways are subject to biochemical noise, which can prevent downstream messengers from serving as an exact reflection of the extracellular signal (52). In the case of CAR-4-1BB signaling, this means that the activation of the canonical NFκB pathway and subsequent changes in gene expression patterns mediated by the nuclear localization of NFκB may not necessarily match the amount of antigen to which the cell is exposed. In order to make assessments about the accuracy of signal transduction of canonical NFκB signaling in CAR-4-1BB cells, we simulated a population of CAR-4-1BB cells in which the accurate population-level response to target antigen was precluded by stochastic variations in the intracellular concentrations of proteins involved in signaling. We were particularly interested in how accurately each successive level of the transduction pathway reflects the amount of antigen stimulation. To this end, we performed simulations by stochastically varying all protein concentrations upstream from a given level of the pathway. We also varied the antigen concentration level. Then, we computed the mutual information between the distribution of responses at this level and the distribution of antigen concentrations (Fig. 5). This allowed us to track how the amount of information carried by the pathway changes at each level of signal transduction. Since experimental measurements on the intrinsic variability of CAR-4-1BB cells were not available, we performed our simulations for multiple values of the noise parameter. In this way, we were able to identify common observable trends that characterize the system, regardless of a certain variability level considered.

**Figure 5:**
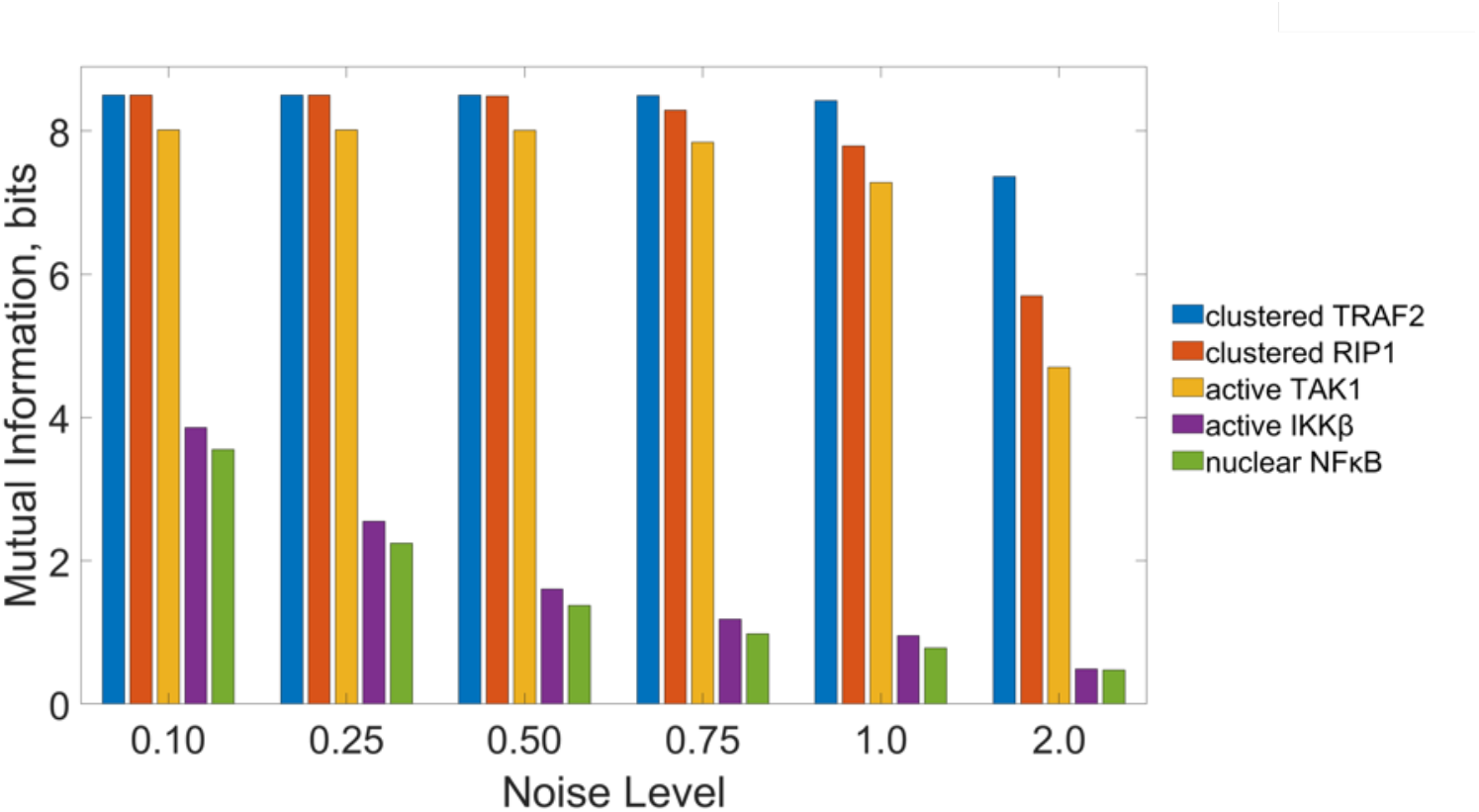
Information transduction along the canonical NFκB pathway. Calculated mutual information in bits between antigen concentration and concentration of clustered TRAF2, clustered RIP1, enzymatically active TAK1, enzymatically active IKKβ and nuclear NFκB.

We can visually observe differences between the distribution of peak NFκB nuclear concentrations does not exactly mirror the antigen distribution (Fig. S1). Specifically, by obtaining the distribution of peak NFκB nuclear concentrations at different intrinsic noise levels, we observed that with increasing noise parameter, the NFκB response shifts towards lower values for the same underlying antigen distribution (Fig. S1).

Our analysis of the mutual information provides a quantitative measure of differences between the antigen distribution and the distribution of concentrations for successive steps in the signaling pathway (Fig. 5). We predict that the proteins recruited early in signal transmission exhibit a relatively small loss of information following antigen stimulation. By definition, the mutual information between two distributions cannot exceed the marginal entropies of each of those distributions (47). The marginal entropy for the distribution of antigen concentrations we used was 8.79, while the mutual information at the level of TARF2 clustering was close to this upper limit, between 7.36 and 8.50 bits, depending on the intrinsic noise level. Interestingly, most information loss consistently occurred between the enzymatic activation of the TAB2/TAK1 complex and enzymatic activation of the NEMO/IKKβ complex. In some conditions, there was up to a 10-fold reduction in the mutual information at this stage. Meanwhile, signal transduction between enzymatically active IKKβ and the nuclear translocation of NFκB seemed relatively accurate with only a small reduction in mutual information. In summary, we find that for all noise levels, mutual information is predicted to decrease as the signal is transduced through the pathway, with the steps directly leading up to the activation of IKKβ as the least accurate step in signal transduction.

### Suggested improvements to CAR design based on mutual information

A primary goal of CAR therapy is to enhance the on-target cytolytic activity of CAR cells while avoiding side effects. Some of these side effects include cytokine release syndrome, on-target-off-tumor effects, and graft-versus-host disease (53). We hypothesized that if secondary messengers more accurately reflect the amount of antigen encountered by the engineered CAR cell, CAR cells would show a more specific response to antigen binding. Particularly, we suggest that increasing the mutual information between the antigen levels and the nuclear concentration of NFκB would affect the therapeutic performance of cells in a manner such that the cells do not become unnecessarily active when encountering only trace amounts of the target, while showing enhanced activity levels when encountering higher amounts of antigen.

Thus, we applied the model to investigate how changing the initial concentration of each protein involved in signal transduction affects the mutual information between antigen concentration and nuclear NFκB concentration (Fig. 6). We repeated our Monte Carlo simulations, this time increasing or decreasing by 50% the initial concentrations for individual proteins. We also considered modulating pairs of proteins that together form multimeric enzymes, such as TAB2/TAK1 and NEMO/IKKβ, to check for possible synergistic effects. The analysis showed that it is possible to increase mutual information by 23% by overexpressing NEMO (Fig. 6A). The overexpression of NEMO yielded the same increase as its joint overexpression with IKKβ, indicating a lack of synergistic effects. Mutual information decreased by 20% when TAB2 was overexpressed separately or in conjunction with TAK1. We believe that these results indicate that overexpressing NEMO experimentally would make the nuclear NFκB concentration in response to activation more reflective of the extracellular antigen concentration. Thus, the potency of the cellular response triggered by NFκB would have a closer correspondence to the targets encountered. When repeating the same procedure for decreased protein concentrations (Fig. 6B), the estimated mutual information between antigen levels and nuclear NFκB is predicted to increase by 43% with TAB2 underexpressed but is reduced by 33% when underexpressing NEMO.

**Figure 6:**
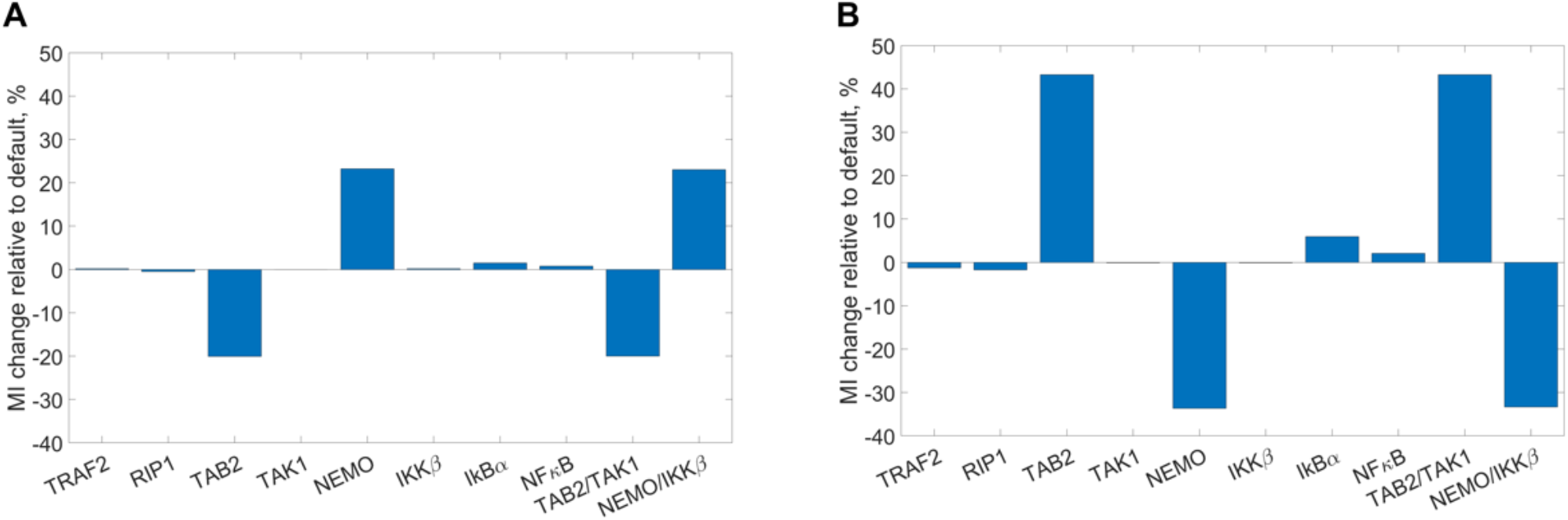
Relative changes in the mutual information. The mutual information between antigen concentration and peak nuclear NFκB is calculated for a noise parameter value of 0.75. (A) Relative change in mutual information due to 50% simulated protein overexpression; (B) Relative change in mutual information due to 50% simulated protein underexpression.

In order to elucidate the mechanism by which the overexpression of NEMO and underexpression of TAB2 cause an increase in mutual information between antigen exposure and NFκB activation, we obtained dose response curves for peak concentrations of active IKKβ (Fig. 7A) and nuclear NFκB (Fig. 7C), and their timings (Fig. 7B, D) in each of those cases. We can see that in cases of intermediate and high antigen concentrations but not low concentrations, the overexpression of NEMO causes an increase in peak concentration attained by active IKKβ, compared to the default concentration levels (Fig. 7A). The enhancing effect of NEMO overexpression on nuclear NFκB peak concentration (Fig. 7C) is limited to intermediate antigen concentrations. Thus, NEMO overexpression alters the response of the canonical NFκB pathway in a way, such that the pathway is slightly more active in case of moderate antigen concentrations without showing such an enhancement at lower antigen concentrations.

**Figure 7:**
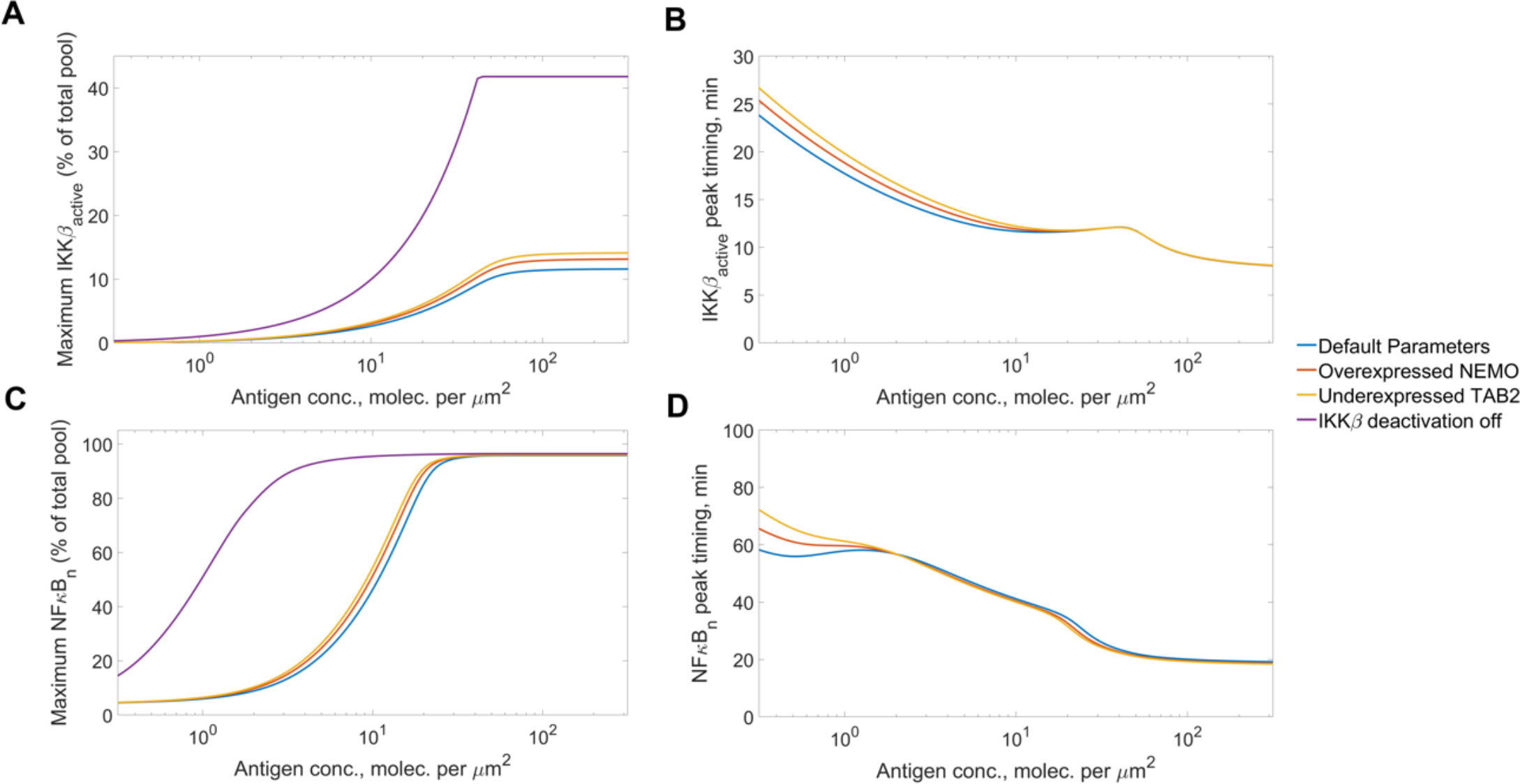
Dose response curves for IKKβ and NFκB in response to antigen binding the CAR with modified expression levels. Peak concentration times and relative magnitudes were recorded for active IKKβ and nuclear NFκB for a range of antigen concentrations and with different proteins under-or overexpressed. (A) Relative magnitude of the peak concentration of active IKKβ; (B) Relative magnitude of the peak concentration of nuclear NFκB; (C) Timing of the peak concentration of active IKKβ; (D) Timing of the peak concentration of nuclear NFκB. N.B. Peak timings for the case of disabled IKKβ deactivation are not shown in (C) and (D) since neither IKKβ nor NFκB show substantial local maxima.

By plotting similar dose response curves for the case of TAB2 underexpression (Fig. 7), we saw an effect by which the decreased amount of TAB2 caused more IKKβ activation. This was highly surprising, since the TAB2/TAK1 complex is known to phosphorylate and activate the NEMO/IKKβ complex, where TAB2 and NEMO are the docking subunits of each multimeric enzyme (36,37). We hypothesized that since both TAB2 and NEMO dock at K63 ubiquitin tails attached to RIP1, the underexpression of TAB2 would make more docking sites available to NEMO and, thus, increase the amount of IKKβ available for activation. Indeed, when we ran simulations to compare concentrations of ubiquitin-bound TAB2 and NEMO, we saw that with underexpressed TAB2, there is a greater abundance of ubiquitin-bound NEMO accompanying the decrease in ubiquitin-bound TAB2 (Fig. S2). This can also explain the greatly reduced mutual information when TAB2 is overexpressed (Fig. 6A), since in this case, TAB2 would outcompete NEMO for access to docking sites. Thus, both over-and underexpression studies underline the centrality of NEMO abundance for enhancing the response of the canonical NFκB pathway.

Finally, given the importance of IKKβ activity for nuclear translocation of canonical NFκB, we investigated how altering the IKKβ activation profile may affect signal transduction. Specifically, we tested whether more persistent IKKβ activity resulting from disabled deactivation would enhance canonical NFκB signaling for low and intermediate antigen concentrations. We were motivated by the fact that such manipulations were shown to be feasible experimentally (39). As expected, the model predicted that lasting IKKβ activation (Fig. S3A) results in more persistent nuclear concentration of NFκB upon antigen stimulation (Fig. S3B).

In order to test the consequences of this hypothetical manipulation on accuracy of signal transduction, we repeated our simulations to compute the mutual information carried by the pathway when IKKβ deactivation is disabled. Notably, the mutual information between antigen concentration and nuclear NFκB concentration showed a dramatic increase for all values of the noise parameter (Fig. S4). In order to find out the reasons for this increase, we plotted dose response curves for the system under disabled IKKβ deactivation (Fig. 7). Deactivation of IKKβ increases the peak level of IKKβ for intermediate to high antigen concentrations (Fig. 7A). On the other hand, there was a substantial amplification of the NFκB response in low and intermediate antigen concentration conditions, with the amplification observed for the intermediate range being much more dramatic (Fig. 7C). This suggests that if IKKβ deactivation is absent, the canonical NFκB pathway would be highly sensitive to antigen concentrations of the order of 1 molecule per µm^2^, without demonstrating as high activation levels for much lower antigen concentrations. Interestingly, the timing of the peak IKKβ or NFκB were not affected by IKKβ deactivation (Fig. 7B, D).

## Discussion

We present a novel mathematical model of NFκB signaling mediated by 4-1BB. Importantly, the model predicts signaling features that agree with what is already known about NFκB signaling. Particularly, prior measurements have shown that IKKβ activity peaks 10-15 minutes after stimulation, and nuclear NFκB peaks approximately 30 minutes after stimulation in the dynamic range of the pathway (54). Thus, the model generates dynamics that have been observed experimentally and can be used to investigate the effects of 4-1BB stimulation.

We applied this model to investigate various characteristics of 4-1BB-mediated signaling. By examining the dose response of active IKKβ and NFκB, we found that the magnitudes of their transient peaks are sensitive to antigen concentration, while the timing of their peaks changes very little within the range of physiologically relevant stimulus concentrations. Next, we used a global sensitivity analysis to identify parameters that have a significant influence on the magnitude of pathway activation. Particularly, our results highlight the influence of NFκB nuclear import/export parameters in determining the peak nuclear concentration of NFκB for most levels of antigen concentrations. Finally, our analysis of mutual information between antigen concentration and successive levels of the NFκB pathway enabled us to suggest specific strategies for enhancing the accuracy of NFκB signaling initiated by 4-1BB. Particularly, overexpression of NEMO and disabling of IKKβ deactivation can greatly increase the information transmission capabilities of the NFκB pathway.

Our modeling analysis provides biologically relevant insight that can be applied to engineered immune cells. CAR-4-1BB Natural Killer (NK) cells are a particularly promising platform for future work. It has been shown that NK-based adoptive cell therapies tend not to suffer from graft-versus-host disease (55–57) and cytokine release syndrome (58–60), disadvantage that frequently accompany CAR-T cell-based therapy (53). Thus, modeling-driven design improvements to CAR-4-1BB NK cells motivated by a greater understanding of cell signaling would contribute to further progress in CAR development.

Prior experimental measurements of signal transduction by the canonical NFκB pathway have shown that the channel capacity (i.e., the maximum possible mutual information) of this pathway is approximately equal to 0.92 bits, which implies that it can resolve only 2^0.92^ ≈ 2 levels of extracellular signal (28). This was interpreted to mean that the pathway is able to only distinguish the presence of the stimulus from its absence. An information theoretic analysis of signaling pathways has shown that even small variations in the number of bits carried by pathways (0.6 bits vs. 0.85 bits) can result in a two-fold difference in the number of cells that exhibit an erroneous response to the extracellular signal (61). Based on this, we suggest that increasing the mutual information carried by the canonical NFκB pathway in response to antigen binding the CAR can result in a closer alignment between CAR-4-1BB cell activity with the antigen concentration it encounters. Thus, our findings can guide further attempts to improve the performance of CAR cells. For example, the predicted improvements can allow the cell to not only to distinguish more accurately whether there is a target or not, but also to discriiminate between various concentrations of the target.

Along with the significant findings produced by our work, we recognize some limitations that can be addressed in the future. One such limitation is that since direct measurements or estimates were not available for some parameter values, we had to make order-of-magnitude estimates by knowing the timescale at which underlying biological phenomena are observed. Values for other kinetic parameters and protein concentrations were taken from multiple sources, all measured or indirectly estimated in fibroblasts with receptors from the TNFR superfamily triggering the activation of NFκB. Our motivation for directly applying these data on CAR-4-1BB is the fact that 4-1BB itself is a member of the TNFR superfamily, and like many other members of the same superfamily, transmits signals by means of TRAF proteins (33,57,62). However, more exact numerical results can be obtained if the model is specifically calibrated for the context of CAR-4-1BB-engineered immune cells. Secondly, even though our model accounts for competition between TAB2 and NEMO for docking sites along K-63 ubiquitin tails, this rests on the assumption that each ubiquitin tail has only two docking sites and that the sites do not discriminate between different docking proteins. While the lack of such competition would likely mitigate the negative effect of TAB2 abundance on NEMO/IKKβ activity, the strong enhancement of NFκB signaling in case of persistent NEMO/IKKβ activation emphasizes NEMO/IKKβ complex’s centrality for enhancing the response of the pathway. Finally, another limitation is the assumed distribution of antigen concentrations that will be encountered by the CAR-4-1BB cells. Since it is known that at least 95% of B-cell acute lymphoblastic leukemia (ALL) cells are CD19-positive (63,64) and that protein concentrations typically follow a lognormal distribution (46), we assumed a lognormal distribution and approximated it to match CD19 antigen levels measured on CD19-positive ALL cells (44). However, specific choices of the antigen distribution could impact our calculation of mutual information. Thus, exploring a broader range of possible distributions would help to form a more complete picture. For instance, the same experimental study had shown that some myeloma cells (termed “CD19-negative”) show vanishingly small CD19 expression levels (around 0.001 molecules per µm^2^) compared to the rest of the population (around 1 molecule per µm^2^). Thus, in the case of myeloma cells, the distribution of CD19 concentrations among the population could be approximated by a bimodal distribution. The percentage of these CD19-negative myeloma cells varied significantly from patient to patient in a range of 10-80% (44). A future study could investigate information transmission capabilities of the NFκB pathway in the context of a bimodal antigen distribution with different percent contributions from each unimodal component.

## Conclusions

Our work focused on developing a mathematical model of CAR-4-1BB-induced NFκB signaling and analyzing its implications. Specifically, we predicted the dose response curves and different activation metrics of the pathway to elucidate the primary features of the signaling dynamics. Using a sensitivity analysis, we identified model parameters that significantly affect the quantitative predictions of our model. Finally, based on an information-theoretic analysis of NFκB signaling, we were able to make suggestions for the further improvement of CAR cell design. Particularly, our results demonstrated that overexpressing NEMO and disabling IKKβ deactivation are predicted to increase the sensitivity of CAR-4-1BB cells against potential therapeutic targets. Our work provides a predictive, quantitative framework that can guide the development and optimization of 4-1BB-induced CAR signaling.

## Abbreviations

ALL: Acute Lymphoblastic Leukemia
CAR: Chimeric Antigen Receptor
eFAST: Extended Fourier Amplitude Test
IκBα: Nuclear factor of kappa light polypeptide gene enhancer in B-cells inhibitor, alpha
IKK: Inhibitor of NFκB Kinase
NEMO: NFκB Essential Modulator
NFκB: Nuclear Factor κB
NK: Natural Killer
RIP1: Ribosome-inactivating Protein
TAB2: Transforming Growth Factor Beta Activated Kinase 1 Binding Protein 2
TAK: Tat-associated Kinase
TRAF2: Tumor Necrosis Factor Receptor Associated Factor 2

## Declarations

### Ethics approval and consent to participate

This article does not represent research with ethics considerations.

### Consent for publication

This article does not use any private or personal data.

## Availability of data and materials

The MATLAB files containing the model used in this work, along with the MATLAB scripts used to perform the analyses presented here are available via GitHub: https://github.com/FinleyLabUSC/CAR-41BB-NFkB-signaling.

## Authors’ contributions

SDF proposed and directed the project. VT constructed the mathematic model, performed simulations and developed the figures. VT wrote the draft manuscript. VT and SDF jointly edited the manuscript. SDF provided the funding.

## Acknowledgements

Both authors would like to acknowledge the members of the Computational Systems Biology group at the University of Southern California for the constructive criticisms and comments.

