## Supplemental Figures for "Computational analysis of 4-1BB-induced NFκB signaling suggests improvements to CAR cell design"

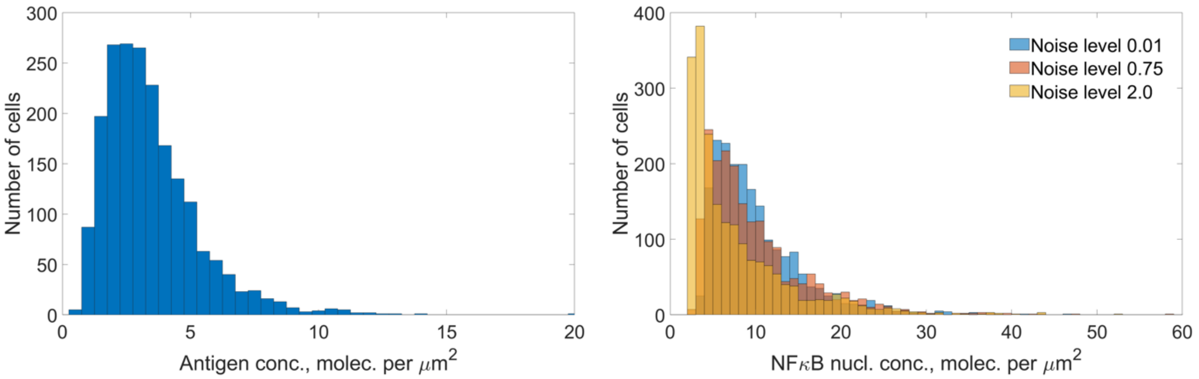


**Figure S1: Distribution of antigen concentrations and peak NFκB nuclear concentrations.** (A) Histogram of the sampled antigen concentration used throughout Monte Carlo simulations; (B) Histogram of peak NFκB nuclear concentrations observed in response to antigen stimulations from (A) at different intrinsic noise levels.

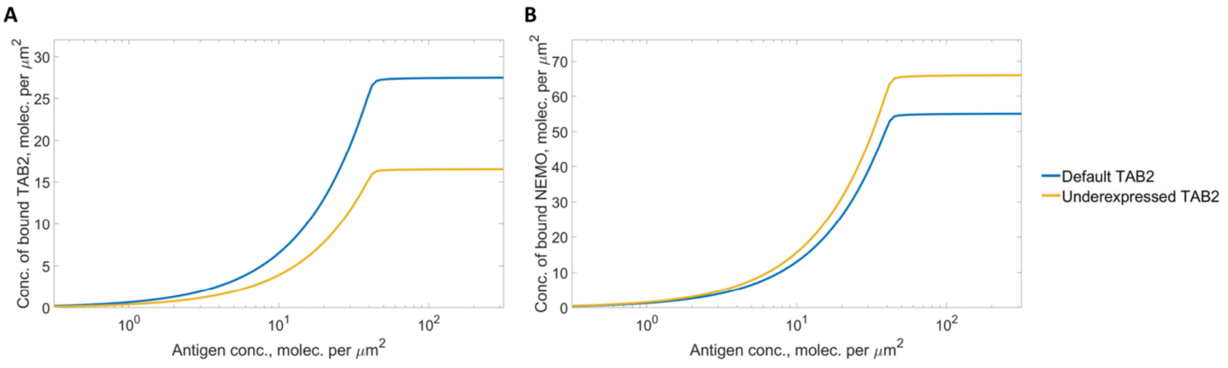


**Figure S2: Dose response curves of K-63 Ubiquitin-bound TAB2 and NEMO with default initial concentration or 50% underexpression of TAB2.** (A) Concentration of bound TAB2 as a function of antigen concentration; (B) Concentration of bound NEMO as a function of antigen concentration.

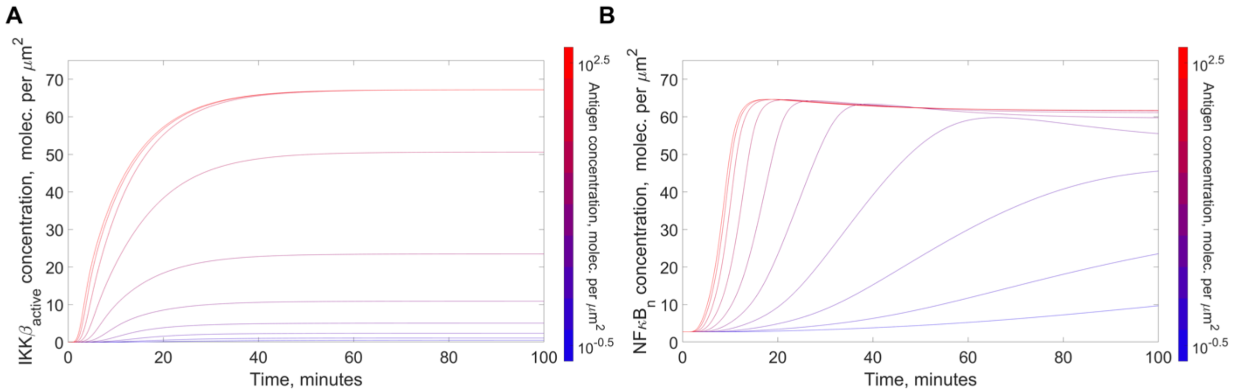


**Figure S3: Activation profiles for IKKβ and NFκB in response to antigen binding the CAR with disabled deactivation of IKKβ.** The pathway was stimulated with 10 different antigen concentrations *in silico* and activation profiles for IKKβ and NFκB were recorded for each antigen concentration. (A) Concentration of enzymatically active IKKβ; (B) Nuclear concentration of NFκB.

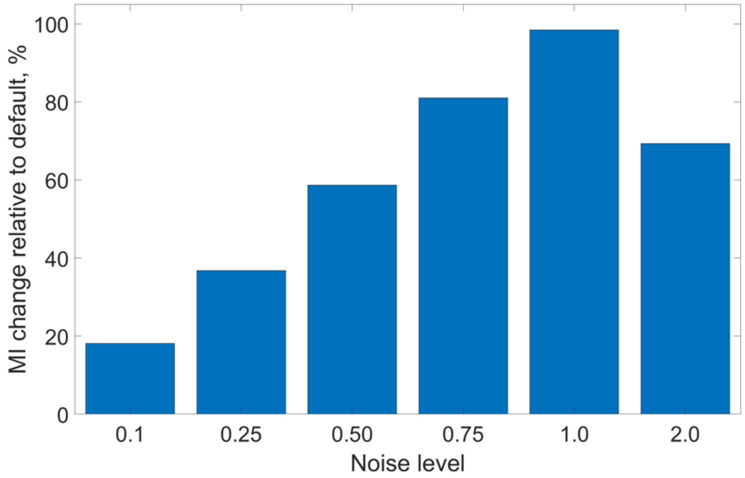


**Figure S4: The transduction of information along the canonical NFκB pathway with disabled IKKβ deactivation.** Percentages show relative decrease of mutual information at the level of nuclear NFκB concentration compared to the default model at different values of the noise parameter.
